# Rapid divergence in independent aspects of the compatibility phenotype in the Spiroplasma/Drosophila interaction

**DOI:** 10.1101/2021.02.03.429608

**Authors:** Joanne S. Griffin, Michael Gerth, Gregory D. D. Hurst

## Abstract

Heritable symbionts represent important components of host biology, both as antagonistic reproductive parasites and as beneficial protective partners. An important component of heritable microbes’ biology is their ability to establish in new host species, a process equivalent to a host shift for an infectiously transmitted parasite or pathogen. For a host shift to occur, the symbiont must be compatible with the host: it must not cause excess pathology, must have good vertical transmission, and possess a drive phenotype that enables spread. Classically, compatibility has been considered a declining function of genetic distance between novel and ancestral host species. Here we investigate the evolutionary lability of compatibility to heritable microbes by comparing the capacity for a symbiont to establish in two novel host species equally related to the ancestral host. Compatibility of the protective *Spiroplasma* from *D. hydei* with *D. simulans* and *D. melanogaster* was tested. The *Spiroplasma* had contrasting compatibility in these two host species. The transinfection showed pathology and low vertical transmission in *D. melanogaster* but was asymptomatic and transmitted with high efficiency in *D. simulans*. These results were not affected by the presence/absence of *Wolbachia* in either of the two species. The pattern of protection was not congruent with that for pathology/transmission, with protection being weaker in the *D. simulans*, the host in which *Spiroplasma* was asymptomatic and transmitted well. Further work indicated pathological interactions occurred in *D. sechellia* and *D. yakuba*, indicating that *D. simulans* was unusual in being able to carry the symbiont without damage. The differing compatibility of the symbiont with these closely related host species emphasises first the rapidity with which host-symbiont compatibility evolves despite compatibility itself not being subject to direct selection, and second the independence of the different components of compatibility (pathology, transmission, protection). This requirement to fit three different independently evolving aspects of compatibility, if commonly observed, is likely to be a major feature limiting the rate of host shifts. Moving forward, the variation between sibling species pairs observed above provides an opportunity to identify the mechanisms behind variable compatibility between closely related host species, which will drive hypotheses as to the evolutionary drivers of compatibility variation.

## Introduction

Heritable symbiont infections – microbes that pass vertically from a female to her progeny -are widespread in nature and exert various effects on insect host reproduction, provide defence against natural enemies and aid in host physiological processes (Moran, 2006). Symbioses can be obligate or facultative, and mutually beneficial as well as costly and parasitic (Buchner, 1965). The impacts of symbionts on their individual hosts has both ecological and evolutionary implications. For instance, symbionts can alter the capacity of their host to colonize new ecological niches or change the direction of sexual selection (Dyson and Hurst, 2004; Salem *et al*, 2020). They may drive host evolution, both to accommodate symbionts with positive effects (for instance, bacteriomes) (Buchner, 1965) and suppress negative symbiont characteristics (Hornett *et al* 2006).

One unusual feature of symbiont-encoded traits is that, in contrast to ‘normal’ genetic traits, heritable microbes can move across species boundaries. Just as a pathogen may undergo a host shift event, causing a spillover event or spreading through a novel species epidemically, so a heritable symbiont may shift to another species and subsequently establish through vertical transmission. Comparison of host and symbiont phylogenies indicates that host shift events occur commonly for some symbionts (Sandstrom *et al*, 2001; Werren *et al*, 1995). It is of particular significance for ‘broad range’ symbionts such as *Wolbachia* and *Spiroplasma* and is also important for some of those with restricted incidence such as *Hamiltonella* within aphids (Moran *et al*, 2005). The capacity to shift to new host species has led to heritable microbe-host evolutionary ecology to be considered conceptually similar to plasmid-microbe dynamics (Jiggins and Hurst, 2011; Dietel *et al*, 2018), and also viewed within the framework of ‘Community Genetics’ (Ferrari and Vavre, 2011). Within this framework, whether a symbiont is common or rare, and the diversity of hosts infected, depends on the rate at which they host shift, and the breadth of hosts they can move into.

For pathogens, compatibility to establish in novel hosts depends at least in part on the relatedness of the recipient to the original host species (Gilbert and Webb, 2007; Streicker *et al*, 2010). For example, pathogenic viruses are more likely to infect host species that are closely related to their ancestral host, such that the success of infection decreases with increasing phylogenetic distance from the ancestral host (Longdon *et al*, 2011). This pattern has also been observed for heritable *Spiroplasma* symbionts in ladybird beetles (Tinsley and Majerus, 2007). However, it has been increasingly recognised that genetic distance is only one component of compatibility, and that closely related host species may share similar levels of susceptibility to a pathogen regardless of their genetic distance from the ancestral host. This ‘phylogenetic effect’ may create variable incidence of infection between different host clades, with some host clades that are distantly related to the microbe’s natural host being compatible with the symbiont (Longdon *et al*, 2014). That compatibility to heritable microbes evolves over time is also indicated by the large evolutionary distances between host species carrying related symbionts.

For a heritable endosymbiont to become established in a novel host population, there must be a route of infection to the novel host and capacity to infect the female germline (Herren *et al*, 2013). Following introduction to the new host, the microbe must then drive into the population. Spread requires an infected female to produce more infected daughters than an uninfected female produces daughters. Compatibility thus has three components. First, there is the vertical transmission efficiency of the symbiont – the rate at which it passes from mother to daughter. Symbionts in novel hosts typically have less faithful vertical transmission than native ones (Clancy and Hoffmann, 1997; Russell and Moran, 2005; Kageyama *et al*, 2006; Tinsley and Majerus, 2007; Nakayama *et al*, 2015). Second, symbionts in novel hosts must not show maladaptive virulence phenotypes (McGraw *et al*, 2002; Carrington *et al*, 2010; Nakayama *et al*, 2015). Third, spread of a symbiont requires the presence of a ‘drive’ in the novel host – either in the form of reproductive parasitism or a direct benefit to the host of carrying the symbiont. This may be in form of retention of previous phenotypes in the novel host, or the appearance of one in the novel interaction.

In this study, we investigate the rate of evolution of host compatibility for *Spiroplasma* in drosophilids. Unlike *Wolbachia*, which resides in the cytoplasm of host cells, *Spiroplasma* occupies a largely extracellular niche in *Drosophila* being commonly found in the haemolymph (Sakaguchi and Poulson, 1961). The relationship between host and *Spiroplasma* can vary from beneficial defensive mutualism to reproductive parasitism (Jaenike *et al*, 2010; Xie *et al*, 2010; Xie *et al* 2014). In *Drosophila hydei, Spiroplasma* protects against the generalist parasitoid wasp, *Leptopilina heterotoma*, without manipulating host reproduction (Xie *et al*, 2010). In its native host it has near perfect vertical transmission efficiency and very low cost of infection (Xie et al. 2009; Corbin, 2017). The *Spiroplasma* in *D. hydei* is related to strains naturally found in willistoni group of flies and *D. melanogaster* (Kageyama *et al*, 2006). Thus, the symbiont can undergo regular host shift events (Haselkorn *et al*, 2009).

The technical ease by which *Spiroplasma* can be experimentally shifted into new hosts was first recognised in the 1960s, when the sex ratio organism (now known to be *S. poulsonii*) was placed into different species, with varying success (Williamson and Poulson, 1979). Hasselkorn *et al*, (2009) exploited this ease of transinfection to investigate the degree to which *Spiroplasma* from *D. neotestacea*, that protects against nematode attack, would thrive in other drosophilids from the mushroom feeding guild that were attacked by the same worm. This study showed varying compatibility for *Spiroplasma*, with just one member of the guild being a compatible host. These studies all presented successes and failure of transinfected *Spiroplasma* but they did not investigate the evolutionary pattern of compatibility.

In this paper, we explored the evolutionary lability of *Spiroplasma* host compatibility by introducing *Spiroplasma* from *D. hydei* into *D. melanogaster* and *D. simulans*. Members of the melanogaster subgroup and *D. hydei* last shared a common ancestor 40-62 MYA (Ranz *et al*, 2001). Previous work indicated that the *Spiroplasma* strain from *D. hydei*, Hy1, misfits in *D. melanogaster* due to poor vertical transmission and virulence in the novel host (Kageyama *et al*, 2006; Nakayama *et al*, 2015). To date, no one has explored the capacity of other species in the *melanogaster* subgroup to propagate Hy1 infection that would indicate the degree to which compatibility is labile. We therefore artificially transferred Hy1 from its native host, *D. hydei*, to the novel hosts, *D. melanogaster, D. simulans* and measured transmission efficiency, fitness costs and protective phenotype against attack by *L. heterotoma*. In addition, we examined whether Hy1 *Spiroplasma* virulence was observed in two other members of the melanogaster subgroup, *D. yakuba* and *D. sechellia*.

## Materials and Methods

### Symbiont and host

*Spiroplasma* infected *D. hydei* (*Spiroplasma* strain TEN 104-106 haplotype 1, Hy1, Mateos *et al*, 2006) was used in this study. Hy1 falls as an out-group of *Spiroplasma poulsoni* and shows no evidence of reproductive manipulation in the *Drosophila* strains used as novel fly hosts (Hutchence, 2011). Two recipient strains of *D. melanogaster* were used, one was *Wolbachia* (*wMelCS*)-infected strain of *D. melanogaster*, Canton S (CS), derived from Montenegro *et al*, 2005 and the other a *Wolbachia* uninfected *D. melanogaster* strain, Oregon-R (see Table 1.). The laboratory populations of *Wolbachia* infected (*wRi*) and uninfected *D. simulans* (F15 and F7, respectively) were derived from iso-female lines supplied by the Centre for Environmental Stress and Adaptation Research, La Trobe University, Australia. The *D. sechellia* line was acquired from a Drosophila Stock Centre in the US. We used *D. sechellia* line 21, carrying the *w*Sh strain of *Wolbachia* and a *Wolbachia* uninfected strain of *D. yabuka*.

**Table 1.** *Drosophila* species and strains used in this study.

| Species | Strain | Wolbachia infection and strain | Source |
| --- | --- | --- | --- |
| *Drosophila melanogaster* | Canton S | Yes, *wMelCS* | Montenegro *et al*, 2005 |
| *Drosophila melanogaster* | Oregon-R | No | Hurst laboratory stock, University of Liverpool |
| *Drosophila simulans* | F15 | Yes, *wRi* | Centre for Environmental Stress and Adaptation Research, La Trobe University, Australia. |
| *Drosophila simulans* | F7 | No | Centre for Environmental Stress and Adaptation Research, La Trobe University, Australia. |
| *Drosophila sechellia* | 21 | Yes, *wSh* | *Drosophila* Stock Centre in the US |
| *Drosophila yakuba* | unknown | No | Ben Longdon, University of Exeter |

*Drosophila hydei, D. melanogaster* and *D. simulans* strains were maintained on a cornmeal-based medium (ASG) consisting of yeast, sugar, maize and nipagin, supplemented with live yeast granules, at 25°C under a 12 h:12 h light:dark cycle with overlapping generations. *D. yabuka* and *D. sechellia* were maintained on malt medium (yeast, malt, maize, nipagin and propionic acid) supplemented with live yeast granules, at 25°C under a 12 h:12 h light:dark cycle with overlapping generations.

### Artificial lateral transfer of Spiroplasma and assessment of vertical transmission

Microinjections were carried out as described by Nakayama *et al*, 2015. In brief, hemolymph was extracted from the thorax of *Spiroplasma* infected *D. hydei* and mixed with sterile PBS. Virgin female *Drosophila* recipients were artificially injected in the abdomen with 0.1-0.2μl of PBS-hemolymph, using a hydraulic positive-pressure microinjection apparatus (Model IM-6, Narushige Ltd, Tokyo, Japan). Injected females were aged for 14 days to let the *Spiroplasma* infection establish in the new host. Each female was then placed with 2 uninfected males of the same *Drosophila* strain*-*to ensure mating success-in a vial containing 15 ml *Drosophila* medium.

Adult injected flies were allowed 4 days to oviposit, after which time the males were discarded and the females frozen at −80°C. All injected females were screened for *Spiroplasma* using a PCR assay. Offspring from unsuccessfully infected females were subsequently used as negative controls. DNA extraction and PCR assays were carried out as described by Nakayama *et al*, 2015. Maternal vertical transmission efficiency was determined as the proportion of infected female progeny of the infected mothers. PCR assays were conducted as follows. Each female was macerated in a 50 μl 5% Chelex (Chelex 100 Resin, Bio-Rad Laboratories, Hercules, CA, USA) solution and 1μl proteinase K, and incubated at 37 °C overnight (Walsh *et al*, 1991). Samples were then heated at 95°C for 10min to denature the proteinase K. PCR amplifications were performed using *Spiroplasma*-specific primers SpoulF (5′-GCTTAACTCCAGTTCGCC-3′) and SpoulR (5′-CCTGTCTCAATGTTAACCTC-3′) as in Montenegro *et al*, 2005. PCR cycling conditions were an initial denature of 1 min 30 s at 94°C, followed by 35 cycles of 15 s at 93 °C, 1min annealing at 47 °C and 1min at 72 °C. The injected females were designated the parental generation.

### Measurement of fitness

An outline of the experiments carried out in this study is provided in Figure 1. Virgin female progeny from both infected and uninfected females (injected in both cases) were collected following eclosion and aged for 10 days. The females were then placed with two uninfected males in small plastic vials (2cm height x 2cm diameter) containing grape jelly agar. Flies were permitted to mate and oviposit for 24 h at 25°C, and then tipped onto fresh agar once a day for further 2 days. Larvae from the first day were discarded. On day 4, the males were discarded, and the adult female *Drosophila* were removed and screened for *Spiroplasma* via PCR assay. Larvae whose grandmothers were infected but mothers tested negative for *Spiroplasma* were discarded. First instar larvae were then picked into vials of fly food at a constant density of 20 larvae per vial, to control for larval conditions. Eclosed virgin females of the F2 generation were collected and aged for 10 days.

**Figure 1.**
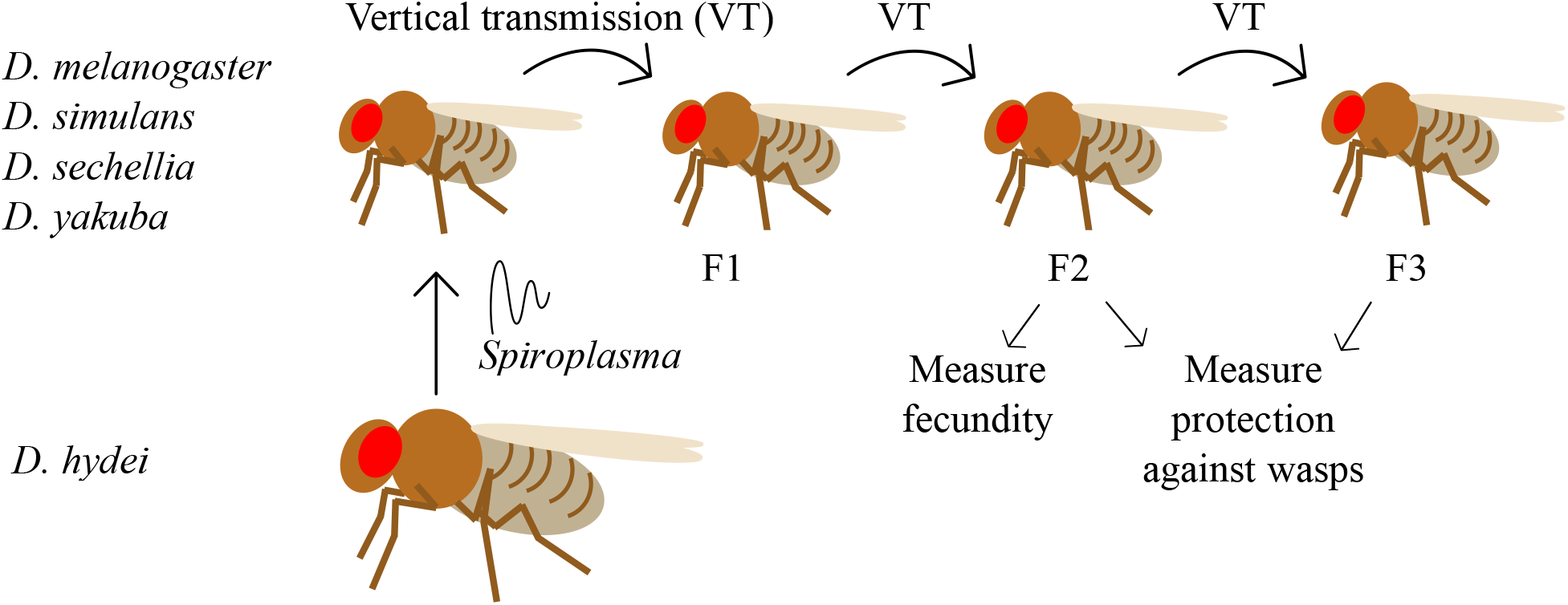
Outline of experimental design. Following lateral transfer of *Spiroplasma* from *D. hydei* to females in the *D. melonagaster* subgroup, females were permitted to mate and oviposit. Following oviposition, females were assayed for *Spiroplasma*. Emerged females of the F1 generation were mated and placed on grape agar jelly to facilitate the collection of larvae. First instar larvae of the F2 generation were picked into vials at a constant density. The number of adult offspring produced by adult F2 females across 4 days was measured. Vertical transmission efficiency was measured at each generation. The protective ability of *Spiroplasma* against parasitoid wasp attack was measured in *D. melanogaster* and *D. simulans* only. Twenty first and second instar *D. melanogaster* and *D. simulans* larvae were exposed to wasps. The number of fly pupae, eclosed adult flies and the number of eclosed wasps were counted for each vial.

To measure the cost of *Spiroplasma* infection in a novel host, we measured the number of adult offspring produced by the F2 generation of infected parental females. Each F2 female was placed with two uninfected males in a vial of fly medium to mate. They were moved into fresh vials every day for five days to allow continuous oviposition. On the sixth day, adult flies were removed, and the females were screened for *Spiroplasma*. Vials that had been occupied by uninfected females (other than the uninfected controls) were kept and noted as flies that had lost infection between the F1 and F2 generations. The eclosed offspring from days 2 to 5 were counted. This provided an index of the direct influence of *Spiroplasma* on host fitness.

We assessed the fecundity of infected females alongside an uninfected control. PCR assays included, a) a positive control re-extracted alongside the test individuals (*D. melanogaster* carrying Melanogaster Sex Ratio Organism: MSRO) and b) repeat PCR assays of negative specimens. Quality control for PCR assays included screening all individuals for the mitochondrially encoded cytochrome c oxidase I (MT-CO1) with samples failing to amplify with this housekeeping marker eliminated.

### Protection assays

We evaluated the ability of *Spiroplasma* to protect its fly host from parasitoid wasp attack following its lateral transfer from *D. hydei* into species in the melanogaster subgroup. We used an inbred laboratory strain of *Leptopilina heterotoma*, originally collected from Sainte Foy-lès-Lyon and la Voulte, France. The strain was donated by Dr Fabrice Vavre and harboured *Wolbachia*. Wasps were maintained on *D. melanogaster* Oregon-R with standard ASG food, at 25°C under a 12 h:12 h light:dark cycle.

The *D. simulans* protection assay was carried out on F3 generation larvae, following artificial infection of adult females with Hy1 (see above). For *D. melanogaster*, F2 generation larvae were exposed to the wasps, following artificial infection with Hy1. For each species, 6 treatments per assay were established: transinfected larvae with and without wasps (S+ Lh+, S+ Lh-), uninfected controls with and without wasps (S-Lh+, S-Lh-) and a positive control naturally infected *D. melanogaster* with and without wasps (MSRO Lh+, MSRO Lh-). For each treatment 10 replicate vials were established, each containing 20 first and second instar *Drosophila* larvae. The larvae were picked onto the surface of standard ASG food from grape agar (as described above). Five mated female *L. heterotoma* adult wasps were immediately added to each wasp treatment vial and left to attack larvae for 3 days at 25°C under a 12 h:12 h light:dark cycle. The wasps had previously been matured to at least 5 days of age at 25°C, with males. The number of fly pupae, eclosed adult flies and the number of eclosed wasps were counted for each vial. This meant that fly and wasp fitness could be assessed in terms of the number of flies and wasps surviving the fly pupal stage.

### Statistical analyses

The number of offspring produced by transinfected and uninfected control females was compared using general linear models. Where there were selection replicates, these were nested within the infection status in the analyses. Where there was only one infected replicate, a paired t-test was used. All statistics was performed using RStudio Software for Statistical Computing, version 1.1.447 (R Core Team, 2013; RStudio Team, 2015).

For the protection assays, the glm() function was used to fit a generalized linear model with a binomial distribution for fly fitness (number of emerging adult flies/number of fly pupae i.e. pupae-to-adult fly survival rate) and wasp fitness (number of emerging adult wasps/number of fly pupae i.e. pupae-to-adult wasp survival rate). The independent variables were *Spiroplasma* infection status and wasp attack (for fly fitness data only).

## Results

### Transmission efficiency and direct cost of Spiroplasma on host fitness in D. simulans and D.melanogaster

*Wolbachia* infected (W+) *D. melanogaster* females infected with *Spiroplasma* produced significantly fewer offspring relative to the uninfected control (Fig.2A, t = −6.217, d.f. = 96, *P* < 0.001). There was no significant effect of replicate (replicate 2, t = 0.079, *P* = 0.937; replicate 3, t = 0.504, *P* = 0.616; replicate 4, t = −0.547, *P* = 0.586, d.f. = 69). *Spiroplasma*-infected W-females also produced fewer offspring relative to an uninfected control (Fig.2B, t = −3.61, d.f. = 87.13, *P* < 0.001). Thus, *Wolbachia* does not affect the susceptibility of *D. melanogaster* to novel *Spiroplasma* infection.

**Figure 2.**
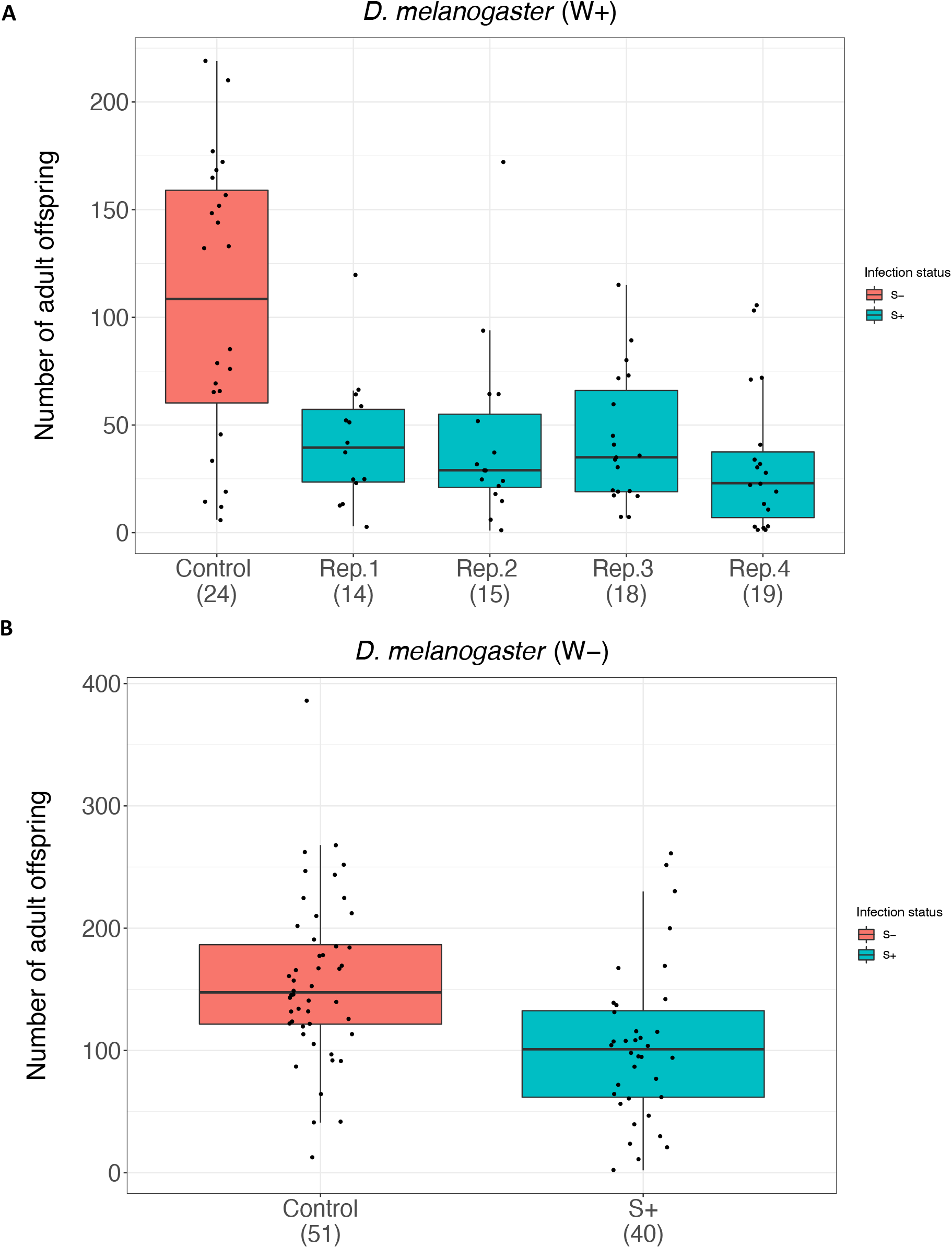
Effect of *Spiroplasma* Hy1 on host fecundity in *D. melanogaster*. The average number of offspring produced over 4 days by the F2 generation of *D. melanogaster* females following artificial lateral transfer of *Spiroplasma* and an uninfected control. **A** *Wolbachia-*infected flies (4 selected infected lines) and **B** *Wolbachia-*uninfected flies (1 selected line). The box plots display the upper and lower quartiles, the median and the range. Points represent each measurement obtained. Values in parentheses show sample size.

In total, 43/69 10-day old *D. melanogaster W*+ F2 females were infected with *Spiroplasma* (62.3%, CI 49.8% −73.7%). The infection prevalence of 75 *D. melanogaster W-*F2 females was 62.7% (CI 50.7% −73.6%).

In contrast, there was no evidence for a difference in the number of offspring produced by *Spiroplasma*-infected *D. simulans* W+ compared to uninfected control individuals (Fig.3A, t = −0.486, d.f. = 74, *P* = 0.628). There was no effect of replicate (replicate 2, t = −1.25, *P* = 0.219; replicate 3, t = 0.848, *P* = 0.400, d.f. = 53). The same was true for *D. simulans* W-(Fig.3B, t = 0.835, d.f. = 91, *P* = 0.406). Again, there was no effect of replicate (replicate 2, d.f. = 73, t = −0.666, *P* = 0.508; replicate 3, t = −1.567, *P* = 0.121).

**Figure 3.**
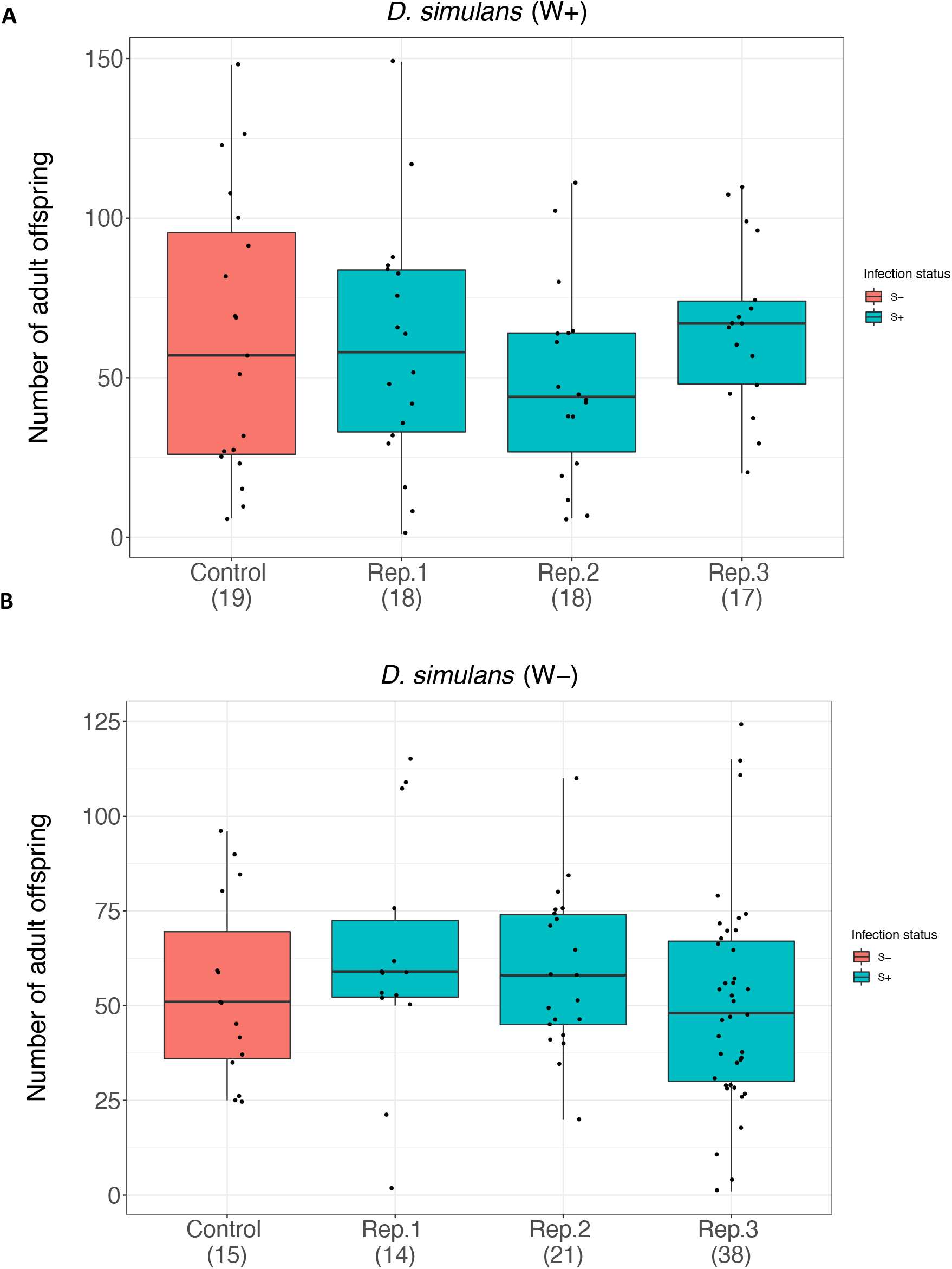
The average number of offspring produced over 4 days by *D. simulans* following artificial lateral transfer of *Spiroplasma* strain Hy1, compared to an uninfected control. **A** *Wolbachia-*infected flies (3 selected infected lines) and **B** *Wolbachia-*uninfected flies (3 selected lines). The box plots display the upper and lower quartiles, the median and the range. Points represent each measurement obtained. Values in parentheses show sample size.

Fifty-six out of 60 F2 generation *D. simulans* W+ were infected with *Spiroplasma* and thus vertical transmission efficiency from the F1 to F2 generation was 93.3% (CI 83.8% −98.2%). Vertical transmission efficiency from the F1 to F2 generation in *D. simulans W*-females was 56.0% (N = 75, CI 44.1% −67.5%).

### Hy1 protection in transinfected Drosophila species

We compared the survival of *Spiroplasma* transinfected- and *Spiroplasma*-free *D. melanogaster* and *D. simulans* in the presence and absence of the parasitoid wasp, *L. heterotoma* (Fig.4A; Fig.5A). *D. melanogaster* naturally infected with the protective *Spiroplasma* strain MSRO was used as a positive control (Fig.S1A and B; Fig.S2A and B). Only results of transinfected flies whose parents had their infection status confirmed by PCR analysis were included.

**Figure 4.**
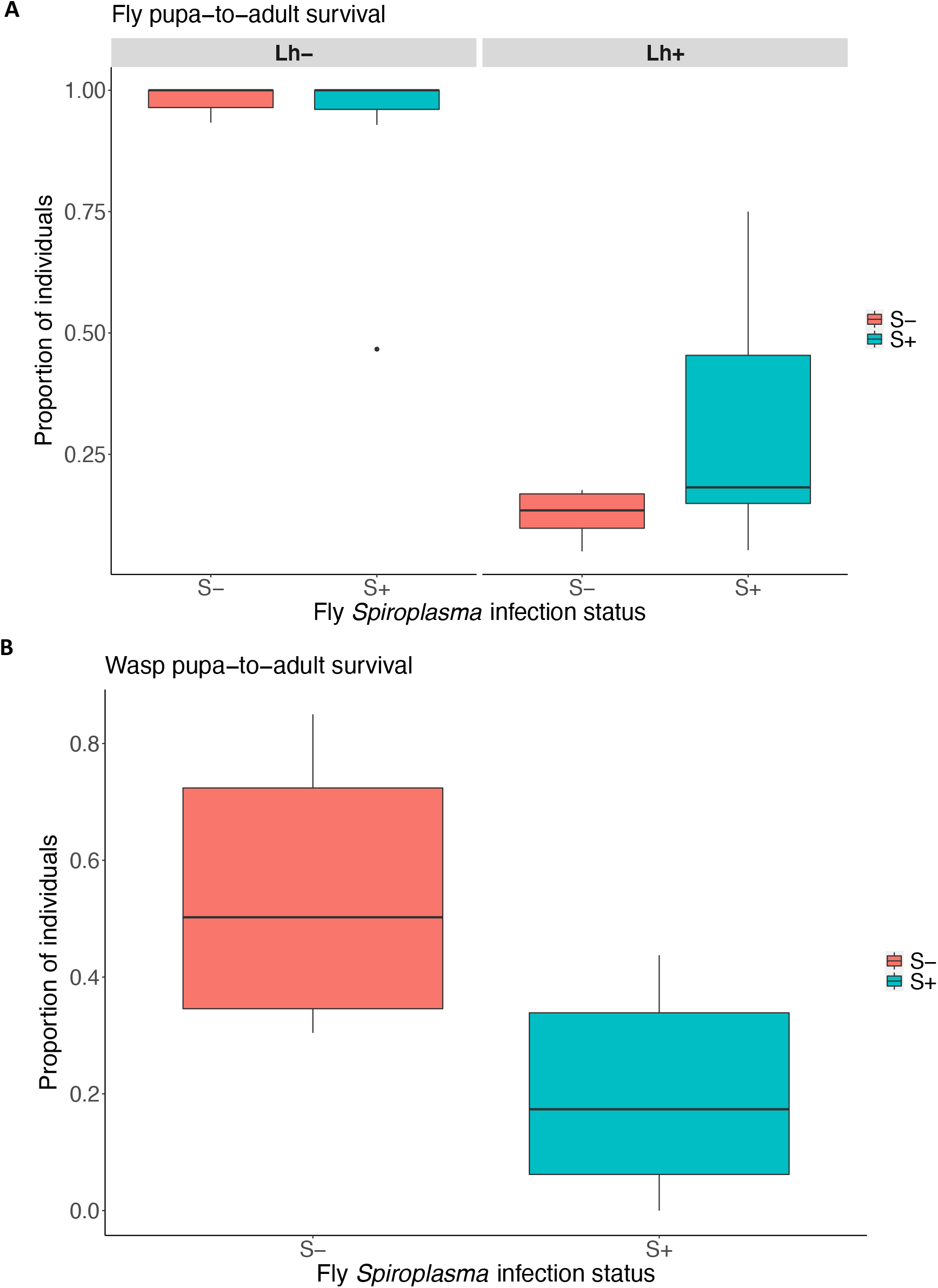
Impact of *Spiroplasma* on wasp parasitism outcome following lateral transfer to *Drosophila melanogaster* W+. **A** Fly pupae-to-adult survival in the presence and absence of *L. heterotoma*. **B** Mean wasp survival in the presence/absence of *Spiroplasma*. Blue box = *Spiroplasma*-transinfected (S+) *D. simulans*; peach box = *Spiroplasma* uninfected (S-) *D. simulans*. The box plots display the upper and lower quartiles, the median and the range.

In the presence of wasps, the pupa-to-adult survival of *Spiroplasma*-transinfected *D. melanogaster* was significantly higher than *Spiroplasma*-free flies (Fig.4A; *z* = 3.35, d.f. = 34, *P* < 0.001). Thus, Hy1 exhibits some degree of protection from wasp-induced mortality in *D. melanogaster*, albeit not to the same extent as MSRO (Fig.S1A). As expected, *D. melanogaster* transinfected with Hy1 gave rise to fewer wasps than *Spiroplasma*-free flies (Fig.4B; *t* = 3.60, d.f. = 12.77, *P* < 0.0033). Wasp pupa-to-adult survival in *Spiroplasma* infected *D. melanogaster* was *c*. 20%, compared to *c*. 54.1% in uninfected flies.

In the absence of wasps, fly pupa-to-adult survival did not differ between *Spiroplasma*-transinfected and *Spiroplasma*-free *D. simulans*. Flies from both treatments had a mean survival of *c*. 100% (Fig.5A). The survival of *Spiroplasma*-transinfected and *Spiroplasma*-free *D. simulans* following attack from *L. heterotoma* was significantly reduced (*c*. 14.3% and *c*. 7.10%, respectively, z = −9.876, d.f. = 35. *P* < 0.001). There was a minor positive effect of *Spiroplasma* on pupa-to-adult survival of *D. simulans* in the presence of *L. heterotoma* but this difference was not significant (z = 0.837, d.f. = 35, *P* = 0.402). Overall, *Spiroplasma* did not significantly enhance the survival of *D. simulans* in the presence of parasitoid wasps. *Spiroplasma*-transinfected flies gave rise to fewer wasps than *Spiroplasma*-free flies (Fig.5B, *c*. 53.7% and *c*. 78.2%, respectively). In other words, wasp pupa-to-adult survival was significantly lower in *Spiroplasma*-transinfected flies (*t* = 3.56, d.f. = 12.3, *P* = 0.0036), thus Hy1 has a negative effect on wasp development without rescuing flies.

**Figure 5.**
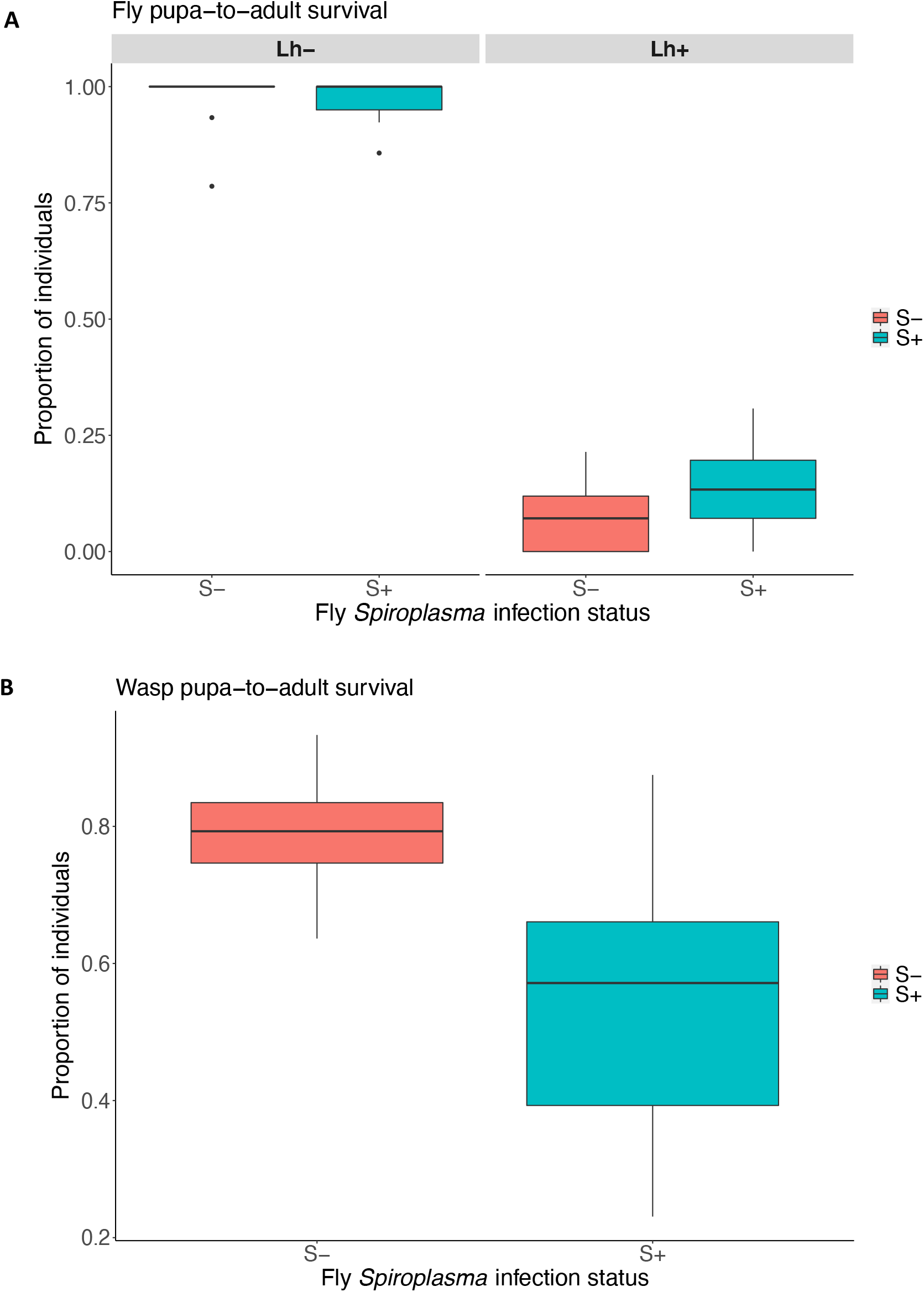
Impact of *Spiroplasma* on wasp parasitism outcome following lateral transfer to *Drosophila simulans* W-. **A** Fly pupa-to-adult survival in the presence and absence of *L. heterotoma*. **B** Average wasp pupa-to-adult survival in the presence/absence of *Spiroplasma*. Blue box = *Spiroplasma*-transinfected (S+) *D. simulans*; peach box = *Spiroplasma* uninfected (S-) *D. simulans*. The box plots display the upper and lower quartiles, the median and the range.

### Impact of Hy1 on D. sechellia and D. yakuba

F2 generation *D. sechellia W+* females infected with *Spiroplasma* produced significantly fewer offspring than uninfected control females (Fig.6. t = −3.52, *P* < 0.001). Vertical transmission efficiency between the F1 and F2 generations was relatively poor, with 36.6% of the F2 maintaining infection (N = 41, CI 22.1% −53.1%). This gave rise to a group of ‘not-infected’ (NI) F2 females, which descended from F1 infected mothers, but tested negative for *Spiroplasma*. The NI group did not produce significantly fewer offspring than the uninfected control (t value = −1.70, *P* = 0.08).

**Figure 6.**
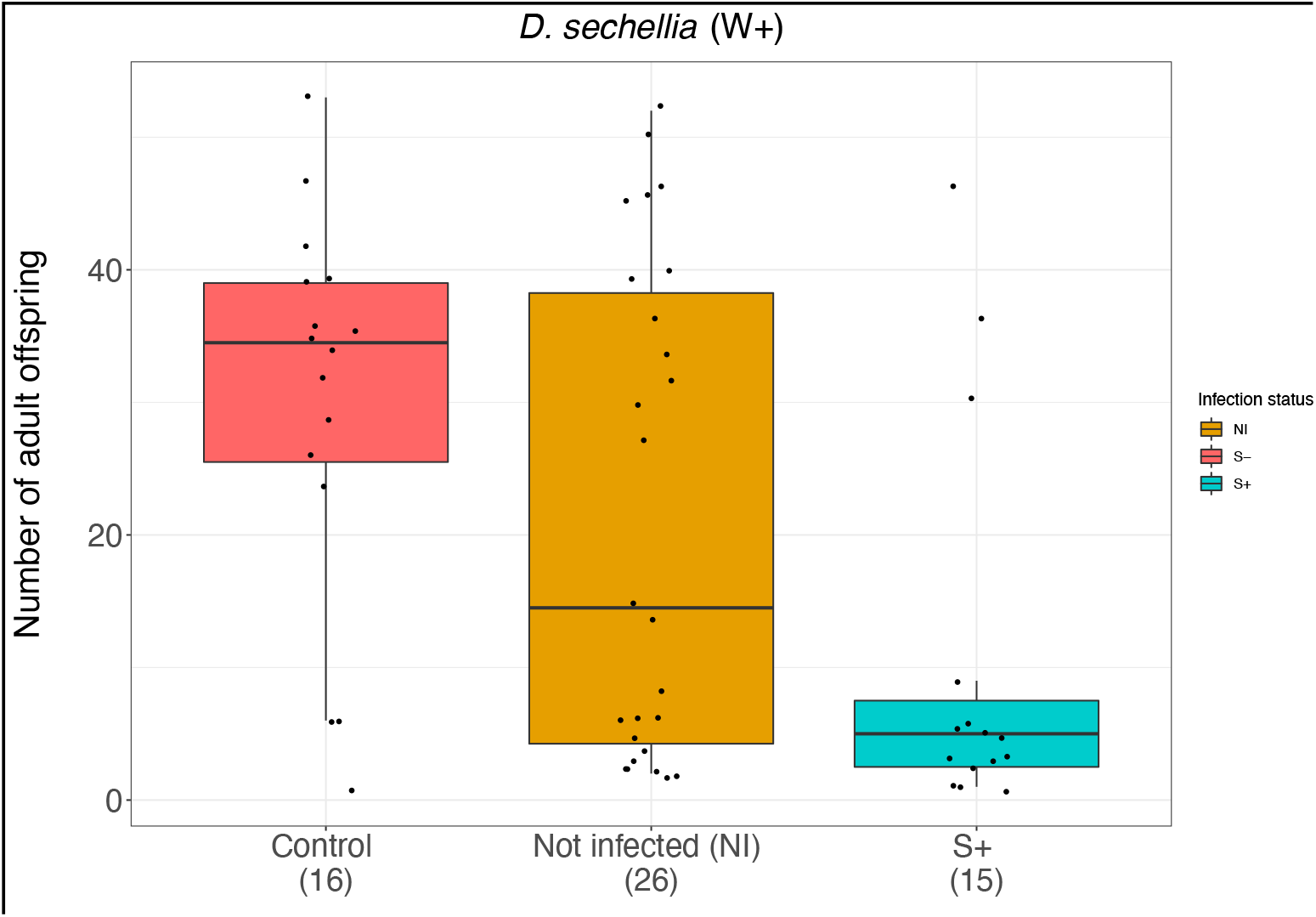
Mean number of offspring produced over 4 days by *D. sechellia W+* following artificial lateral transfer of *Spiroplasma*, and an uninfected control. Females that lost *Spiroplasma* infection between the F1 and F2 generations are called ‘Not infected (NI)’. The box plot displays the upper and lower quartiles, the median and the range. Points represent each measurement obtained. Values in parentheses show sample size.

Four out of 6 *Spiroplasma-*transinfected *D. yakuba W-*lines produced significantly fewer offspring than uninfected controls. Statistically, replicates 5 and 6 did not produce fewer offspring relative to the uninfected control line, but these were characterized by low replication and power (Fig.7, replicate 1, t = −2.63, *P* < 0.05; replicate 2, t = −2.05, *P* < 0.05; replicate 3, t = −2.52, *P* < 0.05; replicate 4, t = −2.39, *P* < 0.05; replicate 5, t = −1.92, *P* = 0.06; replicated 6, t = −1.27, *P* = 0.21). Transmission of *Spiroplasma* between the F2 and F3 generations was exceptionally high, with infection present in 81/83 females (97.6%, CI 91.6% −99.7%).

**Figure 7.**
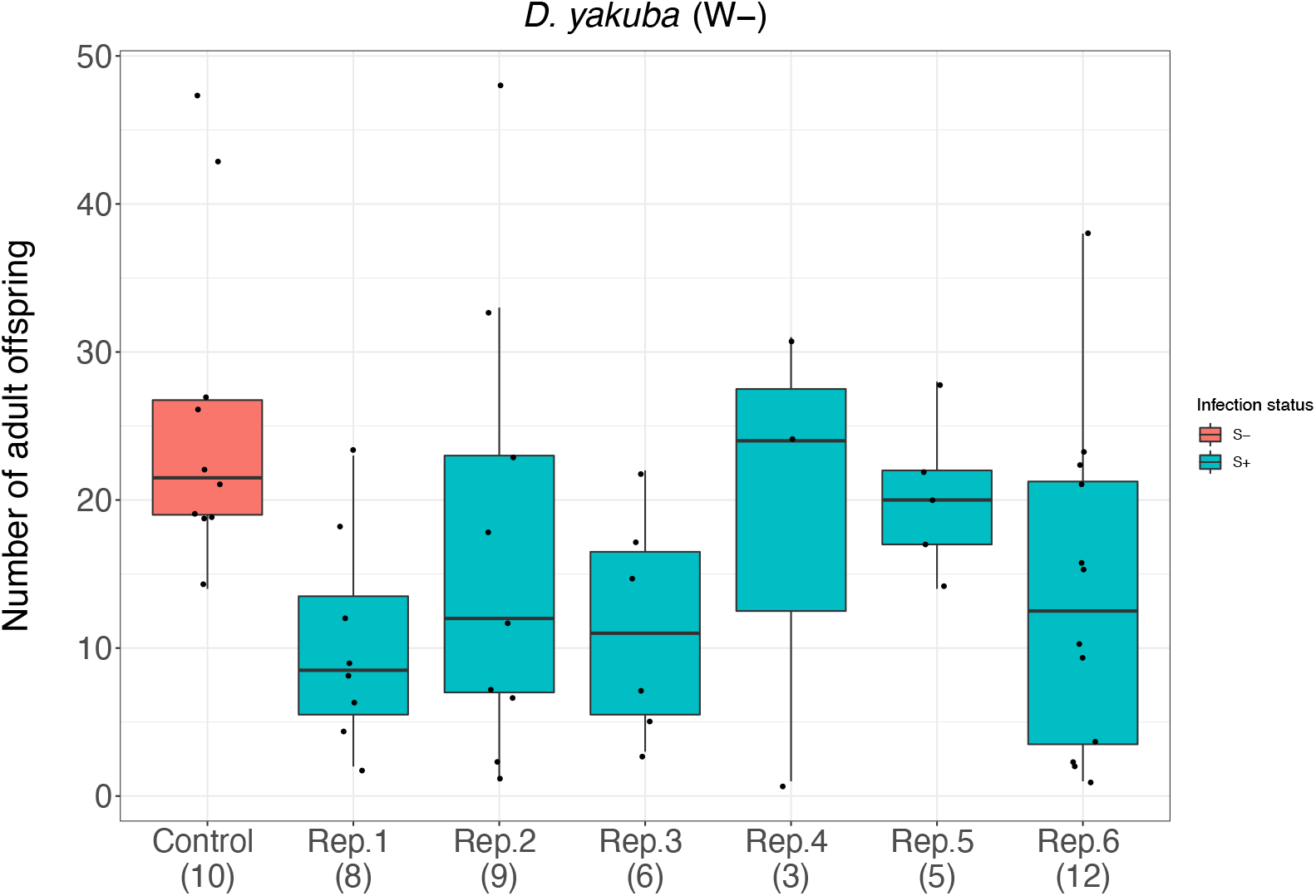
Mean number of offspring produced over 4 days by *D. yakuba W-*following artificial lateral transfer of *Spiroplasma* compared to an uninfected control. The box plot displays the upper and lower quartiles, the median and the range. Points represent each measurement obtained. Values in parentheses show sample size.

## Discussion

The ability of a symbiont to infect and spread in new host species determines the frequency with which host species become infected with symbiont in nature. In this paper we examined one aspect of the host shift process – compatibility with novel hosts. The compatibility of a novel host to a symbiont is a combination of many properties – capacity to vertically transmit, capacity to not cause damage, presence of a phenotype that can drive spread. Our current ‘null’ model of host compatibility holds that the success of a novel infection decreases with increasing distance from the native host. An alternative theory developed for pathogens, is that compatibility evolves across the phylogeny creating a phylogenetic clade effect, in which patches of susceptible hosts are scattered across host phylogeny independently of their distance to the native host (Longdon *et al*, 2014).

We compared host compatibility to novel *Spiroplasma* infection between the closely related species, *D. simulans* and *D. melanogaster*. All species in the *D. melanogaster* subgroup are equally distantly related from *Spiroplasma*’s natural host, *D. hydei*. Similar to Kageyama *et al*, 2006 and Nakayama *et al*, 2015, we find that *Spiroplasma* imposes a direct cost to host fitness and had poor vertical transmission in *D. melanogaster*. This observation was also paralleled in *D. sechellia* and *D. yakuba*. However, *Spiroplasma*-transinfected *D. simulans* did not exhibit pathology relative to the uninfected controls. Vertical transmission was higher in *D. simulans* W+ than in *D. melanogaster* W+ however, vertical transmission was comparable in *D. melanogaster* and *D. simulans* W-individuals. Aside from vertical transmission efficiency, the presence of the secondary endosymbiont, *Wolbachia*, did not influence *Spiroplasma*’s phenotype in the novel host species. It is evident however that Hy1 was more physiologically compatible with *D. simulans* W+ than the other host species tested.

We further examined whether the protective phenotype of *Spiroplasma* could transfer to novel hosts, and if this element of compatibility also diverged between *D. simulans* and *D. melanogaster. Spiroplasma* marginally increased fly survival in *D. simulans* following parasitoid wasp attack and significantly reduced the survival of attacking wasps. In contrast, *Spiroplasma* transinfected into *D. melanogaster* conferred protection, where infected fly hosts showed a significant increase in survival in the presence of parasitoids. The protection in *D. melanogaster* was similar to that observed for the native MSRO infection against this strain of attacking wasp (Jones and Hurst, 2020). *Spiroplasma* induced higher fitness costs in *D. melanogaster* but it better protected the infected individuals from parasitoid attack compared to *D. simulans*. This indicates that the virulence may originate from the protective mechanism. One caveat is that the protection assays were carried out two years after the virulence phenotype observations, such that the strain may have diverged in protection property during laboratory passage.

Together these data indicate that compatibility to *Spiroplasma* Hy1 is evolving within the melanogaster subclade. The nature of this evolution is complex – for pathology and vertical transmission, compatibility is higher in *D. simulans* than in *D. melanogaster*. For the conferred phenotype of protection against natural enemies, compatibility is higher in *D. melanogaster* than in *D. simulans*. Thus, the different aspects of the compatibility phenotype – pathology, vertical transmission, drive phenotype – evolve independently between host species and potentially vary between strains of the same species. The misfit of symbiont to pathology and transmission in one case, and to protection in the other, make it likely that neither would spread in natural populations following introduction. The conditions for spread require alignment of low pathology, high vertical transmission and the presence of a functioning drive phenotype. The sporadic nature of such a multipart alignment, if widely observed, would present a limit to the capacity of this symbiont to spread through the host clade.

Overall, compatibility of closely related hosts to novel *Spiroplasma* infection is an evolutionarily labile trait, in terms of vertical transmission, pathology and protective phenotype. Importantly, species in the *D. simulans* complex diverged only recently. *D. sechellia* and *D. simulans* for example, are estimated to have diverged from one another 413,000 years ago and can produce fertile female hybrids. Nevertheless, these host species differ markedly in their response to novel *Spiroplasma* infection, with pathology observed in one but not the other.

The evolutionary origins of this divergence are unclear. Compatibility, of course, is not a directly selected trait – divergence in compatibility is the product of the evolution of cellular and physiological systems driven by other sources. Our poor knowledge of the mechanistic basis of symbiont-host interactions makes the ultimate source of evolution of compatibility somewhat opaque. It is known that circulating lipid is crucial for *Spiroplasma* success in *Drosophila* (Herren *et al*, 2014), and that vertical transmission is associated with vitellogenin transport into the ovary (Herren and Lemaitre, 2011). Lipid levels, *Spiroplasma* produced RIP toxins, and endogenous defences contribute to protection against wasp attack (Ballinger *et al*, 2017; Kim-Jo *et al*, 2019; Paredes *et al*, 2016). Divergence in these systems would have the indirect consequence of driving changes in compatibility. What is clear is that a model of compatibility based solely on genetic distance of host to symbiont does not explain the complex evolution of this multitrait phenotype. Species that are equally related can react in differing ways to a novel symbiont, and not all aspects of the host-symbiont interaction react in the same way in closely related species.

Finally, it is also notable that compatibility is *Spiroplasma* strain dependent. *Drosophila melanogaster* carries a *Spiroplasma* related to Hy1 in Brazilian and Ugandan populations, and this symbiont shows strong vertical transmission and low pathology. The *D. melanogaster* native *Spiroplasma* strain is most closely related to that found in *Drosophila nebulosa*, a willistoni group drosophilid that is sympatric with *D. hydei* and *D. melanogaster* in South America. Transinfection from *D. nebulosa* to *D. melanogaster* is compatible, with high vertical transmission and low pathology, albeit somewhat a little less stable that the native *Spiroplasma* infection in *D. melanogaster* (Hutchence *et al*, 2012). Whilst these experiments have been conducted in different laboratories and at different times, it is clear that compatibility is additionally evolutionarily labile as a symbiont trait, with quite closely related symbiont strains differing markedly in their compatibility to a given host species. Rapid changes in both symbiont and host determinants of compatibility thus make the outcome of newly introduced strains hard to predict in this system, with neither close relatedness of symbionts, nor close relatedness of hosts, predicting a compatibility.

## Acknowledgments

We thank Jordan Jones for comments on the manuscript, and advice on protection assay design, and Andrea Dewhurst for technical support. JG received funding from Natural Environment Research Council ACCE DTP. Grant Number: NE/L002450/1. This work was supported by the European Commission through H2020 funding in the form of a Marie Curie Fellowship (H2020-MSCA-IF-2015, 703379) to MG.

## Author contributions

Experiments were designed by JG, MG and GH. Experiments were carried out by JG and MG. Data was analysed by JG and MG with advice from GH. Writing. The manuscript was written by JG and edited by GH and MG. All authors approved the final version of the manuscript.

## Data Availability

Raw data underpinning this project can be accessed at

https://doi.org/10.6084/m9.figshare.c.5287780.

## References

1. Ballinger, M.J. Perlman, S.J. (2017) Generality of toxins in defensive symbiosis: Ribosome-inactivating proteins and defense against parasitic wasps in Drosophila. PLoS Pathogens. 13(7): 1–19. https://doi.org/10.1371/journal.ppat.1006431

2. Buchner, P. (1965) Endosymbiosis of Animals with Plant Microorganisms. New York: Interscience.

3. Carrington, L.B. Hoffmann, A.A. Weeks, A.R. (2010) Monitoring long-term evolutionary changes following Wolbachia introduction into a novel host: the Wolbachia popcorn infection in Drosophila simulans. Proc. R. Soc. Lond. B. 277:2059–2068

4. Clancy, D.J. Hoffman, A.A. (1997) Behaviour of Wolbachia endosymbionts from Drosophila simulans in Drosophila serrata, a novel host. Am Nat. 149:975–988

5. Corbin, C. (2018) The evolutionary ecology of an insect-bacterial mutualism. PhD thesis, University of Liverpool.

6. Dietel, A.K. Kaltenpoth, M. Kost, C. (2018) Convergent Evolution in Intracellular Elements: Plasmids as Model Endosymbionts. Trends Microbiol. 26(9):755–768. doi:10.1016/j.tim.2018.03.004.

7. Dyson, E. A. Hurst, G. D. D. (2004) Persistence of an extreme sex-ratio bias in a natural population Proc. Natl. Acad. Sci. USA 101:6520–23

8. Ferrari J, Vavre F. (2011) Bacterial symbionts in insects or the story of communities affecting communities. Philos Trans R Soc Lond B Biol Sci. 12;366(1569):1389–400. doi:10.1098/rstb.2010.0226. PMID: 21444313; PMCID: PMC3081568.

9. Gilbert, G.S. Webb, C.O. (2007) Phylogenetic signal in plant pathogen-host range. Proc. Natl. Acad. Sci. 104(12): 4979–4983. https://doi.org/10.1073/pnas.0607968104

10. Haselkorn, T.S. Markow, T.A. Moran, N.A. (2009) Multiple introductions of the Spiroplasma bacterial endosymbiont into Drosophila. Molecular Ecology. 18(6): 1294–1305. https://doi.org/10.1111/j.1365-294X.2009.04085.x

11. Herren, J.K. Paredes, J.C. Schüpfer, F. Lemaitre, B. (2013) Vertical Transmission of a Drosophila Endosymbiont Via Cooption of the Yolk Transport and Internalization Machinery. MBio. 4(2): 1–8. https://doi.org/10.1128/mbio.00532-12

12. Hornett EA, Charlat S, Duplouy AMR, Davies N, Roderick GK, Wedell N, et al. (2006) Evolution of Male-Killer Suppression in a Natural Population. PLoS Biol. 4(9): e283. https://doi.org/10.1371/journal.pbio.0040283

13. Hutchence, K.J. Fischer, B. Paterson, S. Hurst, G.D.D. (2011) How do insects react to novel inherited symbionts? A microarray analysis of Drosophila melanogaster response to the presence of natural and introduced Spiroplasma. Molecular Ecology. 20(5): 950–958. https://doi.org/10.1111/j.1365-294X.2010.04974.x

14. Hutchence, K.J. Padé, R. Swift, H.L. Bennett, D. Hurst, G.D. (2012) Pheno-type and transmission efficiency of artificial and natural male-killing Spiroplasma infections in Drosophila melanogaster. J. Invertebr. Pathol. 109: 243–247.

15. Jaenike, J. Unckless, R. Cockburn, S.N. Boelio, L.M. Perlman, S.J. (2010) Adaptation via symbiosis: recent spread of a Drosophila defensive symbiont. Science. 9;329(5988): 212–5. doi:10.1126/science.1188235

16. Jiggins, F. M. Hurst, G. D. D. (2011) Rapid insect evolution by symbiont transfer. Science. 332, 185–186.

17. Jones, J.E. Hurst, G.D.D. (2020) Symbiont-mediated protection varies with wasp genotype in the Drosophila melanogaster–Spiroplasma interaction. Heredity 124: 592–602.

18. Kageyama, D. Anbutsu, H. Watada, M. Hosokawa, T. Shimada, M. Fukatsu, T. (2006) Prevalence of a non-male-killing Spiroplasma in natural populations of Drosophila hydei. Applied and Environmental Microbiology. 72(10): 6667–6673. https://doi.org/10.1128/AEM.00803-06

19. Kim-Jo, C. Gatti, J. L. Poirié, M. (2019) Drosophila cellular immunity against parasitoid wasps: A complex and time-dependent process. Front. Physiol. doi:10.3389/fphys.2019.00603.

20. Longdon, B. Hadfield, J.D. Webster, C.L. Obbard, D.J. Jiggins, F.M. (2011) Host phylogeny determines viral persistence and replication in novel hosts. PLoS Pathogens. 7(9): e1002260. https://doi.org/10.1371/journal.ppat.1002260

21. Longdon, B. Brockhurst, M.A. Russell, C.A. Welch, J.J. Jiggins, F.M. (2014) The Evolution and Genetics of Virus Host Shifts. PLoS Pathogens. 10(11) https://doi.org/10.1371/journal.ppat.1004395

22. Mateos, M. Castrezana, S.J. Nankivell, B.J. Estes, A.M. Markow, T.A. Moran, N.A. (2006) Heritable Endosymbionts of Drosophila. Genetics. 174(1): 363–376. https://doi.org/10.1534/genetics.106.058818

23. McGraw, E.A. Merritt, D.J. Droller, J.N. O’Neill, S.L. (2002) Wolbachia density and virulence attenuation after transfer into a novel host. Proc. Natl. Acad. Sci. 99:2918–2923

24. Moran, N.A. Russell, J.A. Koga, R. Fukatsu, T. (2005) Evolutionary relationships of three new species of Enterobacteriaceae living as symbionts of aphids and other insects. Appl Environ Microbiol. 71(6):3302–10. doi:10.1128/AEM.71.6.3302-3310.2005. PMID: 15933033; PMCID: PMC1151865.

25. Moran, N.A. (2006) Symbiosis. Curr Biol. 16(20):R866–71. doi:10.1016/j.cub.2006.09.019. PMID: 17055966.

26. Nakayama, S. Parratt, S.R. Hutchence, K.J. Lewis, Z. Price, T.A.R. Hurst, G.D.D. (2015) Can maternally inherited endosymbionts adapt to a novel host? Direct costs of Spiroplasma infection, but not vertical transmission efficiency, evolve rapidly after horizontal transfer into D. melanogaster. Heredity. 114(6): 539–543. https://doi.org/10.1038/hdy.2014.112

27. Paredes, J.C. Herren, J.K. Schüpfer, F. Lemaitre, B. (2016) The Role of Lipid Competition for Endosymbiont-Mediated Protection against Parasitoid Wasps in Drosophila. MBio. 7(4): 1–8. https://doi.org/10.1128/mbio.01006-16

28. Ranz, J.M. Casals, F. Ruiz, A. (2001) How malleable is the eukaryotic genome? Extreme rate of chromosomal rearrangement in the genus Drosophila. Genome Res. 11:230–239.

29. Russell, J.A. and Moran, N.A. (2005) Horizontal Transfer of Bacterial Symbionts: Heritability and Fitness Effects in a Novel Aphid Host. Applied and Environmental Microbiology. 71:7987– 7994.

30. Sakaguchi, B. Poulson, D.F. (1961) Distribution of “sex-ratio” agent in tissues of Drosophila willistoni. Genetics. 46(12):1665–1676

31. Salem, H. Kirsch, R. Pauchet, Y. Berasategui, A. Fukumori, K. Moriyama, M. Cripps, M. Windsor, D. Fukatsu, T. Gerardo, N. M. Symbiont digestive range reflects host plant breadth in herbivorous beetles. Curr. Biol. 30 (2020), pp. 2875–2886

32. Sandstrom, J.P. Russel, J.A. White, J.P. Moran, N.A. (2001) Independent origins and horizontal transfer of bacterial symbionts of aphids. Molecular Ecology. 10, 217–228.

33. Streicker, D.G. Turmelle, A.S. Vonhof, M.J. Kuzmin, I.V. McCracken, G.F. et al. (2010) Host phylogeny constrains cross-species emergence and establishment of rabies virus in bats. Science. 329: 676–679.

34. Tinsley, M.C. Majerus, M.E.N. (2007) Small steps or giant leaps for male-killers? Phylogenetic constraints to male-killer host shifts. BMC Evol. Biol. 7: 238.

35. Walsh, P.S. Metzger, D.A. Higuchi, R. (1991) Chelex 100 as a medium for simple extraction of DNA for PCR-based typing from forensic material. Biotechniques. 10(4):506–513.

36. Werren, J.H. Zhang, W. Guo, L.R. (1995) Evolution and phylogeny of Wolbachia: Reproductive parasites of arthropods. Proc Biol Sci. 2601(1360): 55–63.

37. Xie, J. Vilchez, I. Mateos, M. (2010) Spiroplasma bacteria enhance survival of Drosophila hydei attacked by the parasitic wasp Leptopilina heterotoma. PLoS ONE. 5(8): e12149. https://doi.org/10.1371/journal.pone.0012149

38. Xie, J. Butler, S. Sanchez, G. Mateos, M. (2014) Male killing Spiroplasma protects Drosophila melanogaster against two parasitoid wasps. Heredity. 112(4):399–408. https://doi.org/10.1038/hdy.2013.118

