## Supplementary Information for "Rapid divergence in independent aspects of the compatibility phenotype in the Spiroplasma/Drosophila interaction"

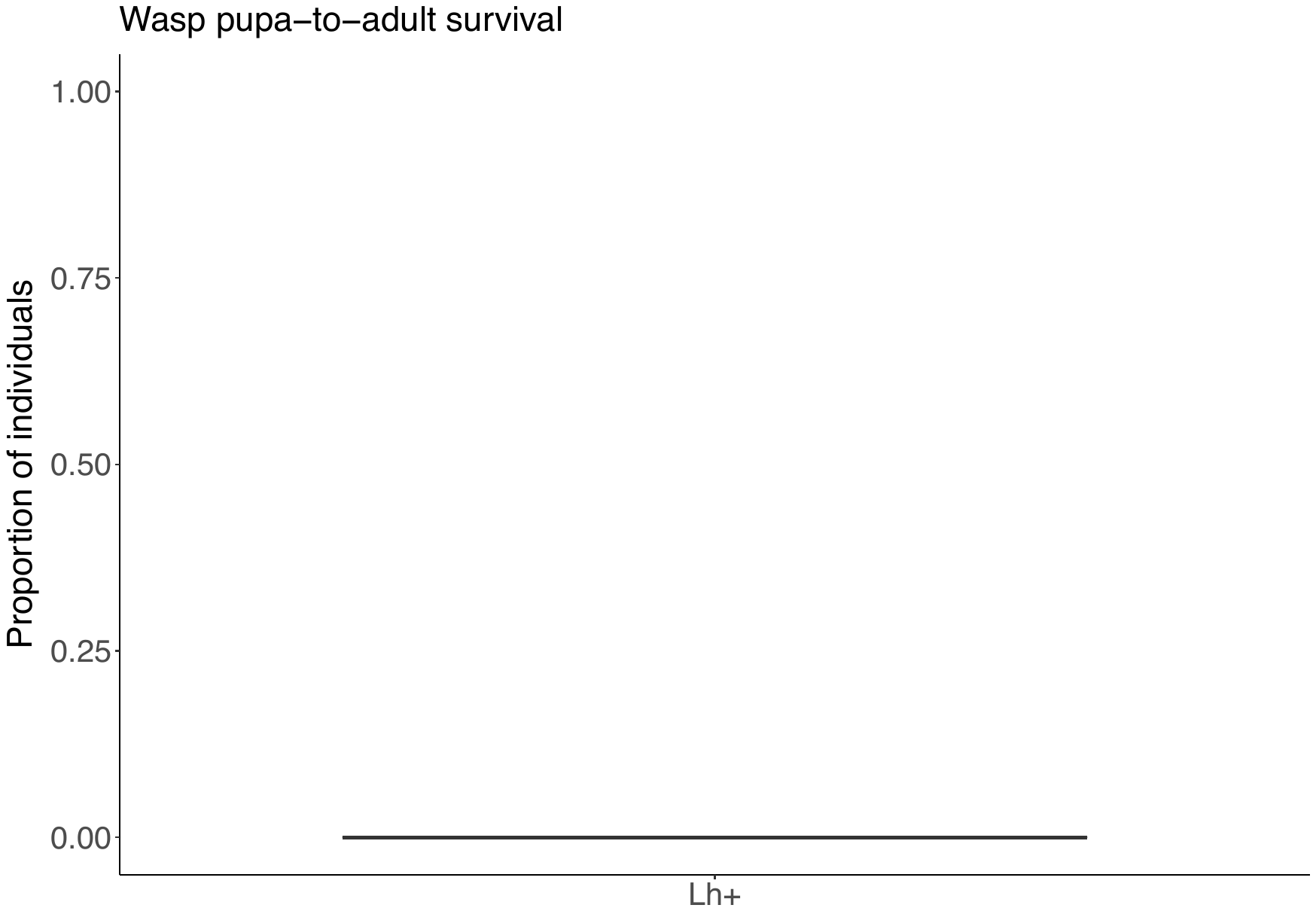

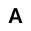

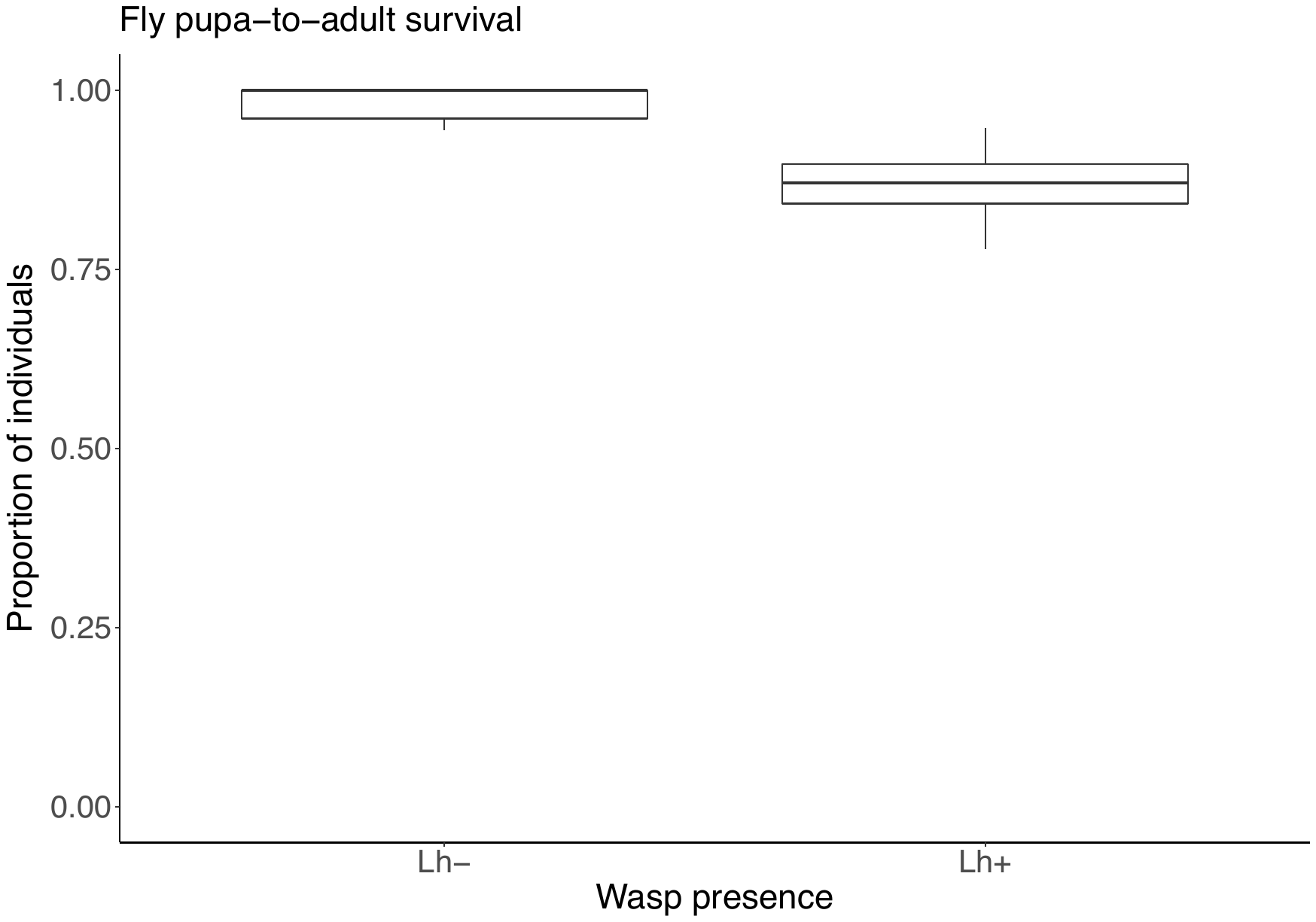

**B**

**Figure S1, relating to Figure 4.** Positive control. Impact of natural *Spiroplasma* MSRO infection on *Drosophila melanogaster* in the presence and absence of wasps. **A** The proportion of flies that emerged in the presence and absence of *L. heterotoma*. **B** Proportion of emerging wasps following wasp attack. The box plots display the upper and lower quartiles, the median and the range.

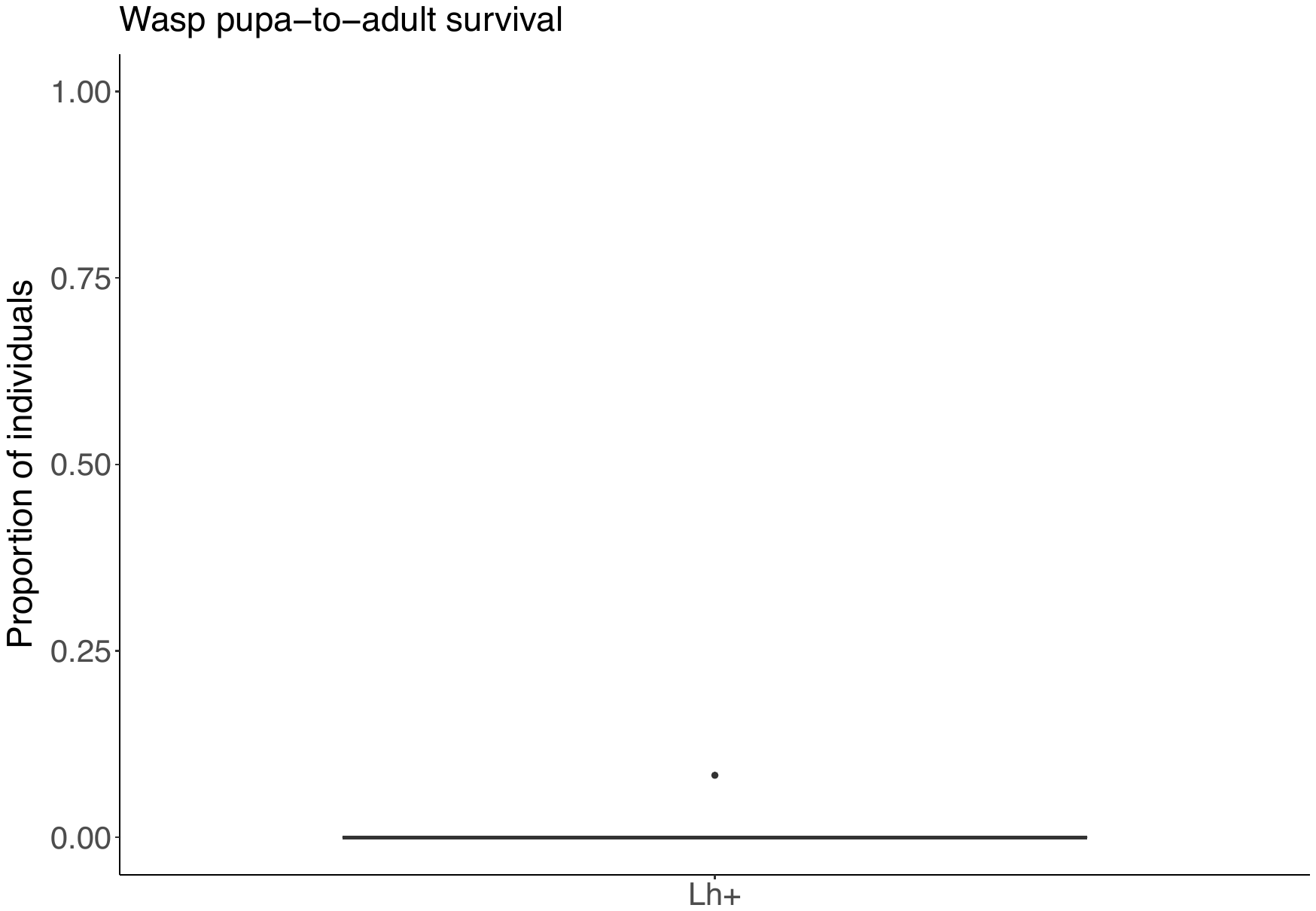

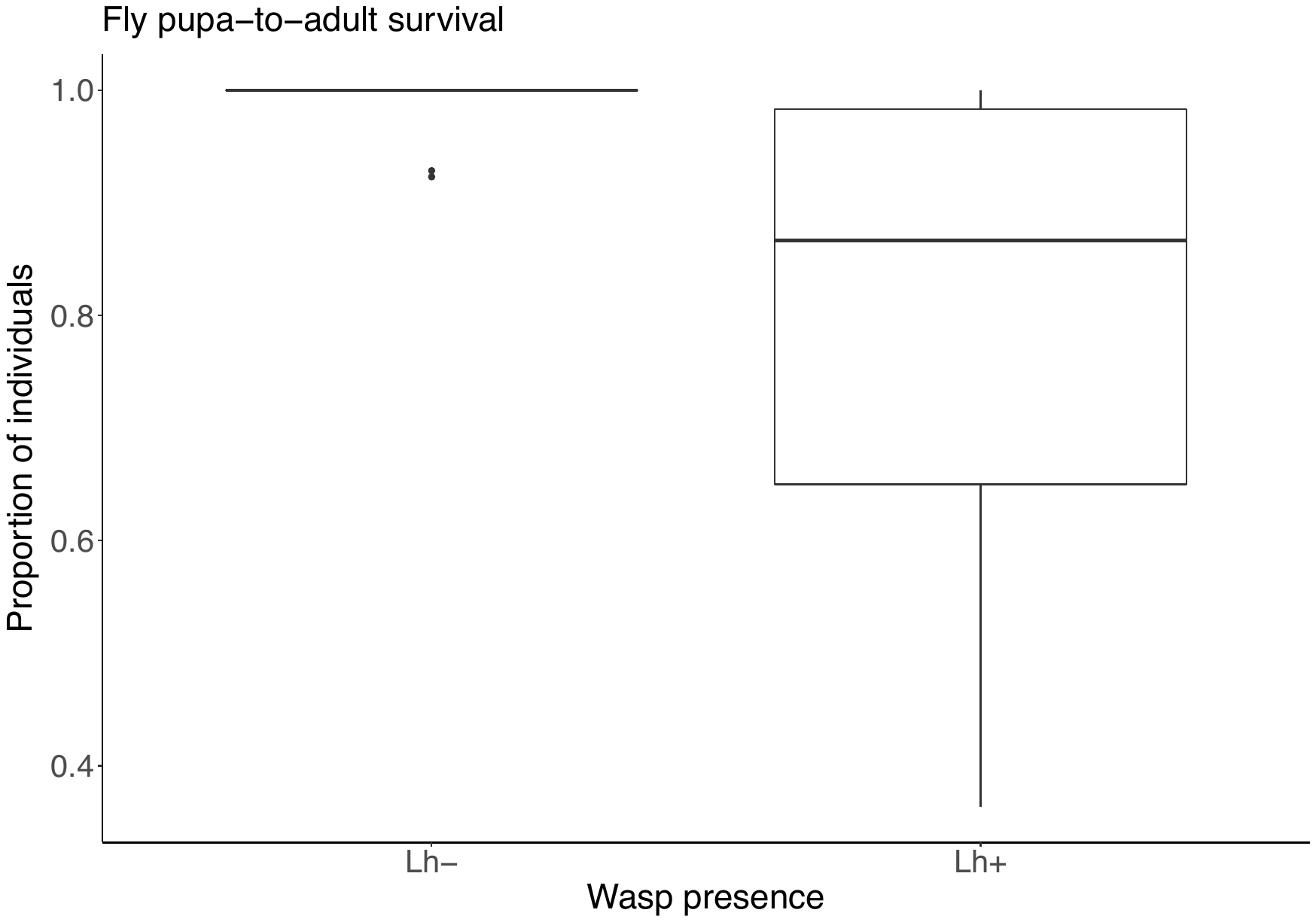

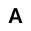

**B**

**Figure S2, relating to Figure 5.** Positive control. Impact of natural *Spiroplasma* MSRO infection on *Drosophila melanogaster* in the presence and absence of wasps. **A** The proportion of flies that emerged in the presence and absence of *L. heterotoma*. **B** Proportion of emerging wasps following wasp attack. The box plots display the upper and lower quartiles, the median and the range.
